# *μ*-PBWT: Enabling the Storage and Use of UK Biobank Data on a Commodity Laptop

**DOI:** 10.1101/2023.02.15.528658

**Authors:** Davide Cozzi, Massimiliano Rossi, Simone Rubinacci, Dominik Köppl, Christina Boucher, Paola Bonizzoni

## Abstract

**Motivation:** The positional Burrows-Wheeler Transform (`PBWT`) has been introduced as a key data structure for indexing haplotype sequences with the main purpose of finding maximal haplotype matches in *h* sequences containing *w* variation sites in 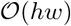-time with a significant improvement over classical quadratic time approaches. However the original PBWT data structure does not allow queries over the modern biobank panels of haplotypes consisting of several millions of haplotypes, as they must be kept entirely in memory.

**Results:** In this paper, we present a method for constructing the run-length encoded PBWT for memory efficient haplotype matching. We implement our method, which we refer to as *μ*-`PBWT`, and evaluate it on datasets of 1000 Genome Project and UK Biobank data. Our experiments demonstrate that the *μ*-`PBWT` reduces the memory usage up to a factor of 25 compared to the best current PBWT-based indexing. In particular, *μ*-`PBWT` produces an index that stores high-coverage whole genome sequencing data of chromosome 20 in half the space of its BCF file. In addition, *μ*-`PBWT` is able to index a dataset with 2 million haplotypes and 2.3 million sites in 4 GB of space, which can be uploaded in 20 seconds on a commodity laptop. *μ*-`PBWT` is an adaptation of techniques for the run-length compressed BWT for the PBWT (RLPBWT) and it is based on keeping in memory only a small representation of the RLPBWT that still allows the efficient computation of set maximal matches (SMEMs) over the original panel.

**Availability:** Our implementation is open source and available at https://github.com/dlcgold/muPBWT. The binary is available at https://bioconda.github.io/recipes/mupbwt/README.html

**Contact:** Paola Bonizzoni

## 1 Introduction

Improved haplotype phasing in large cohorts is facilitating the comprehensive collection and study of variations at chromosome-level for genome evolution and clinical applications. This has been demonstrated by the haplotype-resolved whole-genome sequence data collected from hundreds of thousands of individuals for projects such as the UK Biobank [1] and TOPMed projects [2]. In the field of phased genomics, the positional Burrows-Wheeler Transform (PBWT), which is a data structure that represents permutations of each column of a *h* × *w* binary matrix `M`[1 ..*h*][1 ..*w*], is a key instrument, offering compact representation and efficient haplotype matching for large haplotypes datasets [3]. Indeed, due to the intrinsic capability of the PBWT of saving space in memorizing haplotype data and even in analyzing large haplotypes panels, it is becoming a relevant data structure also in the field of computational pangenomics [4].

Although the PBWT is a vital solution for analyzing pangenomic haplotype data, it has not received as much attention as the famous Burrows-Wheeler Transform (BWT). The seminal paper on BWT by Burrows and Wheeler has been cited more than 3500+ times, which is almost two orders of magnitude larger than that for first PBWT paper by Durbin (364 times acc. Google Scholar). As a result, efficient construction and representation of the PBWT on large datasets is in a relatively nascent stage by comparison to the `BWT`. As a matter of fact, analysis of data such as the UK Biobank data remains to be challenging. In November 2022, Jared Simpson tweeted: *What is the largest publicly available haplotype reference panel? 1000 genomes? I’m looking for a pre-built PBWT index but don’t want to go through dbGAP to get the HRC panel*. Unfortunately, there are not yet solutions to this tweet as the only response was “*We used UK biobank a lot. But it’s also behind the door*.” by Deghi Zhi. The underlying question remains then as to how to efficiently build a PBWT index in an efficient manner that can be used on a commodity machine.

One of the main goals of the original work proposing the PBWT data structure, was to develop a means to find maximal haplotype matches in a set of *h* sequences, each containing *w* variation sites and represented in a matrix `M`. The main idea behind the construction of the PBWT is that of stably sorting the rows of `M` in co-lexicographic order (i.e., sorted order from right-to-left). Durbin [3] showed that maximal haplotype matches can be found in *O*(*hw*)-time since it amounts to finding set-maximal exact matches (SMEMs) using the PBWT, where a SMEM is defined to be the longest fully-terminated match covering a positionbetween the pattern and the string. In this context, it is assumed that each variation site is bi-allelic, meaning that there exists only two observed alleles at a locus in the genome and no insertions or deletions. Although this binary encoding of genetic information appears to remove significant information, it is common practice in the analysis of variations of diploid species, where variations are filtered to only contain bi-allelic sites [5, 6].

Since its initial development, the PBWT has been applied and extended in numerous ways. It has been used for genotype imputation [7], and to create a genotype database search method that is privacy-preserving (PBWT-sec) [8]. Novak et al. [9] and Sirén et al. [10] used the PBWT to encode a graph for haplotype matching (g-PBWT) and graph pangenome indexing [4]. Sanaullah et al. [11] replaced all arrays with linked lists to define a dynamic version of the PBWT (d-PBWT). The original `PBWT` has been used to compute all-pairs Hamming distances [12] and for finding all maximal perfect haplotype blocks in linear time [13].

In this paper, we consider the problem of Durbin [3] that aims to find SMEMs in haplotype data using the PBWT. We demonstrate how it can be efficiently constructed and stored in run-length encoded space. Run-length encoding is a concept that was originally motivated and applied to the BWT; if you consider the BWT for large repetitive input then it is witnessed that there are long repetitions of the same character, which are referred to as *runs*. The number of runs is routinely denoted as *r*, where *r* is usually significantly smaller than *n* on repetitive input. Hence, Mäkinen and Navarro [14] noticed that the the BWT can stored in 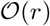 space while still efficiently supporting some standard queries (i.e., count). Although this was noticed by Durbin in 2014, run-length encoding has not been explored since then. Here, we implement a run-length encoding of the `PBWT`, which we refer to as *μ*-`PBWT`, and compare it to Durbin’s PBWT [3] and Syllable-PBWT [15] on both 1000 Genome Project [16] data and simulated panels [1], achieving improvements in the space needed to store the PBWT index.

We demonstrate that *μ*-`PBWT` uses from 1.1 to 25 time less space than Syllabe-PBWT, while uses up to 25000 times less space than Durbin’s PBWT at the cost of up to 2× increase in construction and query time. The experiments show that the best performance of *μ*-`PBWT` is achieved on whole genome sequences data (UK-Biobank chromosome 20 and simulated date). We showed a proof of concept of the scalability of *μ*-`PBWT` to today’s biobanks by producing an index of 13GB for high-coverage whole genome sequencing data on chromosome 20 (stored in a 29GB BCF file). Finally, we show that *μ*-`PBWT` is able to store and query a dataset with 2 million haplotypes and 2.3 million sites in 4 GB of space, which could be easily store on a commodity laptop.

## 2 Preliminaries

### 2.1 Positional Burrows-Wheeler Transform

We define a sequence *S* over a finite, ordered alphabet Σ = {*c*_1_, …, *c_σ_*} of *σ* characters to be the concatenation of *n* characters *S* = *S*[1..*n*]. We denote the empty sequence as *ε*. We denote the *i*-th prefix of *S* as *S*[1..*i*], the *i*-th suffix as *S*[i..*n*], and the sequence spanning position *i* through *j* as *S*[i..*j*], with *S*[i..*j*] = *ε* if *i* > *j*.

The Positional Burrows-Wheeler Transform has been introduced by Durbin as a data structure for handling a matrix `M`, representing a set *S* = {*S*_1_, …, *S_h_*} of *h* sequences of length *w* and over a binary alphabet, simply called haplotypes, by updating two arrays for each column *j*: the *prefix array* `PA`_*j*_ and the *divergence array* `DA`_*j*_.

1. `PA`_*j*_ is the ordering of {1,…, *h*} induced by the co-lexicograph ordering of prefixes of *S* up to column *j* – 1, i.e. formally `PA`_*j*_[*i*] = *k*, if *S_k_*[1..*j* – 1] is the *i*-th element in co-lexicographically ordered list of prefixes *S*_1_[1..*j* – 1], …, *S_h_*[1..*j* – 1].
2. `DA`_*j*_[*i*] stores the length of the longest common suffix between the sequences of index `PA`_*j*_[*i*] and `PA`_*j*_[*i* – 1] up to the (*j* – 1)-th column.

The PBWT of `M` is another matrix `PBWT`[1..*h*][1..*w*] that has the first column identical to the one of `M` while the *j*-th column of `M` with *j* > 1 is obtained by stably sorting the rows of `M` [1 ..*h*][1 ..*j* – 1] in co-lexicographic order. We note that we denote the `PBWT` matrix as `PBWT`. Assuming to denote the *j*-th column of a matrix `M` by `col`(`M`)_*j*_, formally `col`(`PBWT`)_1_ = `col`(`M`)_1_ and `col`(`PBWT`)_*j*_[*i*] = `col`(`M`)_*j*_[`PA`_*j*_[*i*]] for all *i* = *1*..*h* and *j* = 2..*w*.

The main idea is that the prefix-array stores in each column j the permutation of the rows induced by a co-lexicographic ordering of the previous columns up to column *j* – 1 while the divergence array stores in column *j* and position *i* the length of a longest common suffix between row i and the previous one in the permutation induced by the prefix array in column j. Together these two arrays allow to efficiently compute matching queries over haplotype sequences. We note that we frequently use *n* = *h* · *w* to bound the space- and time-complexity.

If we consider the PBWT shown in Figure 1 and Column 5, then `DA`[5][7] = 3 because the co-lexicographically 6-th and 7-th row-prefixes (corresponding to PA[5][6] = 18 and PA[5][7] = 16 rows in the input matrix) up to Column 4 are 0100 and 1100 and their longest common suffix 100 has length 3.

**Figure 1:**
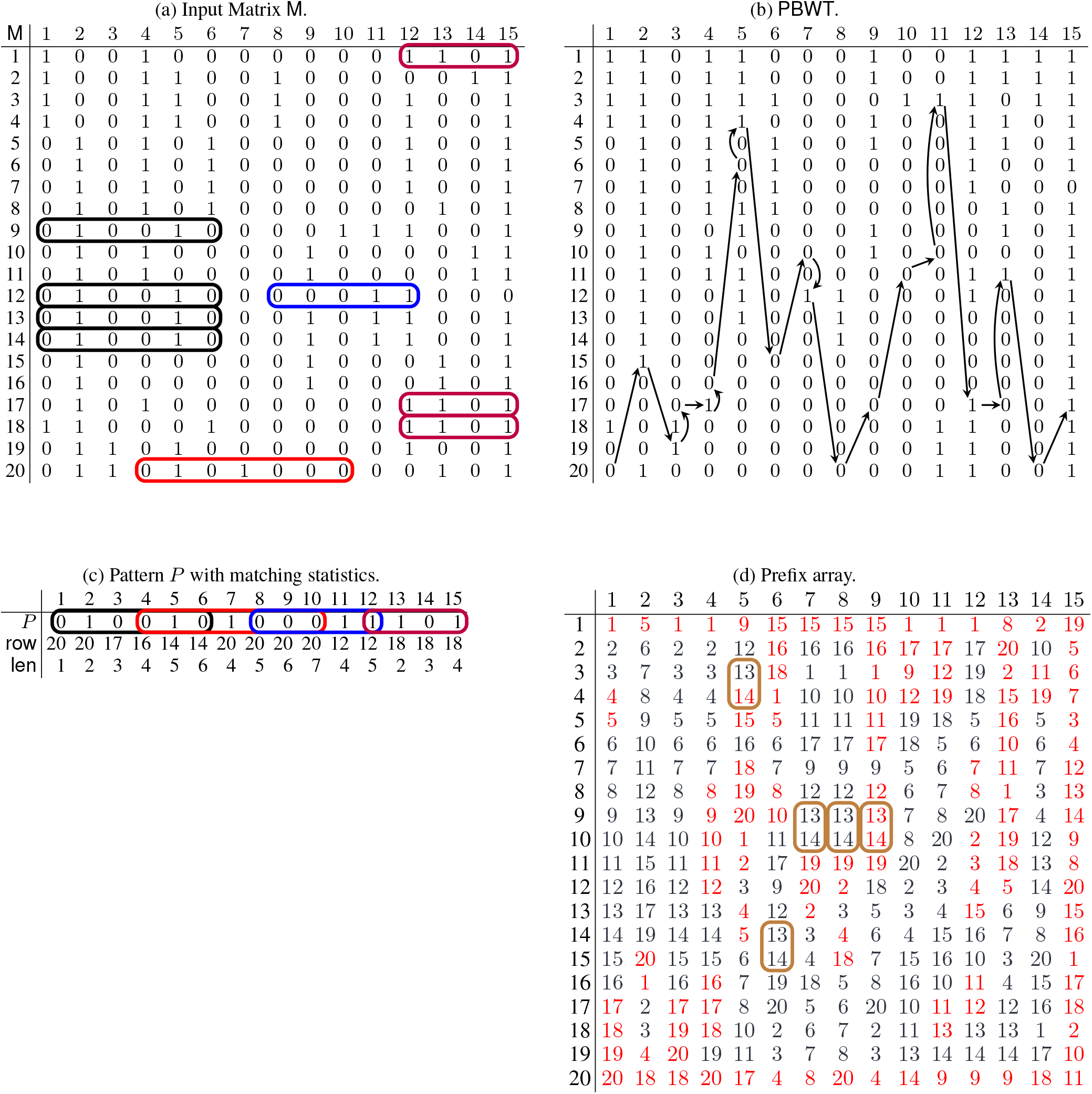
The input matrix `M` of 20 individuals of 15 bi-allelic sites (a), its `PBWT` (b), a query pattern *P* and its matching statistics with respect to `M` (c), the prefix array (PA) of `M` (d). SMEMs are circled in both the pattern and the input matrix `M`. We illustrate the SMEM-finding via computation of matching statistics. We start from an arbitrary row. In this case, we choose the 20-th, where `col`(`PBWT`)_1_[20] = 0. Since we have that *P*[1] = 0, we proceed to the next column, and store `A`[1].`row` = 20 and `A`[1].`len` = 1. To advance by column, we compute the mapping function of row 20 from first column to the second. Observe that the mapping function is used to compute index *k* of `col`(`PBWT`)_*i*+1_ that contains *A*[_*i*_].`row` (details in section 3). Hence, the result is that we are mapping to `col`(`PBWT`)_2_[15]. At the second column, we have *P*[2] = `col`(`PBWT`)_2_[15], so we can proceed to the next column, storing `A`[2].`row` = 20 and `A`[2].`len` = 2. Following the mapping of the row 20, we move onto `col`(`PBWT`)_3_[19]. We have a mismatch at column 3 since *P*[3] ≠ `col`(`PBWT`)_3_[19]. At this point, we can move to either the last character of the previous run, `col`(`PBWT`)_3_[17], or the first character of the next run, `col`(`PBWT`)_3_[20], having `PA`_3_[17] = 17 and `PA`_3_[20] = 18. If we look at the input matrix `M`, we have that, up to column 3 excluded, row 17 has a common suffix to row 20 longer than row 18. So, the best option to maximize the length of the current match is to move to row 17, storing `A`[3].`row` = 17 and `A`[3] .len = 3. Now we use row 17 to compute the mapping function from column 3 to column 4. We proceed in this way until we complete the computation of `A`. Finally, with a sweep from left to right over `A`, we can compute all the SMEMs looking at the indices where `A`[*i*].`len` ≥ `A`[*i* + 1].`len`, as shown through the colored rounded boxes covering *P*.

### 2.2 Run-Length Encoded PBWT

Durbin noted that run-length encoding—originally described by Makinen et al. [14]) for the BWT—can be adapted to the PBWT. We denote the run-length encoded PBWT matrix as `RLPBWT`. This extension is made by observing that the the concept of *run* can be defined for the `PBWT`, i.e., the number of runs in the `PBWT` as the number of binary substrings containing occurrences of the same symbol which are maximal in length. Given *r_j_* as the number of runs in a `RLPBWT` column, we denote *r* as 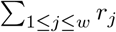. In the following we will use the term PBWT without a specific distinction with the RLPBWT, as the RLPBWT distinguishes for the components it uses.

### 2.3 Set-Maximal Exact Matches

One of the fundamental tasks of the `PBWT` is one-vs-all set-maximal exact matches (SMEMs) finding: the main idea is finding the longest common matching substrings between an external sequence *P* and any other sequence of the same length that are represented in the `PBWT`. Formally, given *w*-length input sequences *S* = {*S*_1_, …, *S_h_*} (sorted in `M`) and a pattern *P*[1..*w*], we define *P*[i..*j*], where 1 ≤ *i* ≤ *j* ≤ *w*, to be a SMEM if it occurs in one of the input sequences of *S* and one of the following holds: i) *i* = 1 and *j* = w; ii) *i* = 1 and *P*[1..*j* + 1] does not occur in *S*; iii) *j* = *w* and *P*[*i* – 1..*w*] does not occur in *S*; iv) *P*[*i* – 1..*j*] and *P*[i.. *j* + 1] do not occur in *S*.

We next define two problems related to finding the SMEMs. First we define the problem of identifying the SMEMs in the pattern *P*.

#### Problem 1 (SMEM-finding)

*Given a set S* ={ *S*_1_, …, *S_h_*} *of h sequences of length w and a pattern P*[1..*w*], *find the list L of pairs* (*p, ℓ*) *such that for all* (*p, ℓ*) ∈ *L*, *P*[*p*..*p* + 1 *ℓ* − 1] *are the SMEMs between S and P*.

Then we define the problem of locating all the occurrences of the SMEMs in the panel.

#### Problem 2 (SMEM-locating)

*Given a set S* ={*S*_1_,…, *S_h_*} *of h sequences of length w and a pattern P* [1..*w*], *find the list L of triples* (*p, ℓ, O*) *such that for all* (*p, ℓ, O*) ∈ *L, P* [*p*..*p* + *ℓ* − 1] *is an SMEMs between S and P where O is the list of haplotypes where the SMEM occur*.

Durbin’s Algorithm 5 [3] is able to solve Problem 2 in 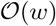-time and 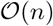-space, which corresponds to about 13n bytes. This memory consumption is the major downside of this algorithm and the motivation that led us to develop a run-length encoded `PBWT` that supports SMEMs finding and locating. For example, in Figure 1 (a), we have 9 SMEMs computed by the pattern *P* in Figure 1 (c).

## 3 Methods

Our main contribution is a significant reduction in the memory used to store the `PBWT` via efficient sampling and storing the PA and DA arrays. In particular, we reduce the space of Durbin’s `PBWT`, which is 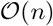-space, to 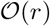-space. And while most of the PBWT operations which require 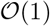-time in Durbin’s `PBWT`—which explicitly stores the input matrix and the associated divergence and prefix arrays—take 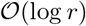-time, this runtime is not observed in practice. We point to the experimental result for illustration of this fact in section 4. Lastly, we refer the reader to Bonizzoni et al. [17] for a more thorough evaluation of the data structures for the PBWT that support different time/space tread-offs for SMEM-finding.

### 3.1 Overview of *μ*-PBWT

The problem of finding SMEMs can be cast into the problem of computing matching statistics for *P*. Given a pattern *P*[1..*w*], the matching statistics of *P* with respect to *S* are an array *A*[1..*w*] of (`row, len`) pairs such that for each position 1 ≤ *j* ≤ *w A*[*j*].`row` is one row of the input matrix `M` where a longest shared common suffix of length *A*[*j*].`len`, ending in position *j* in the pattern *P* and in *S*_*A*[*i*.row_, occurs. SMEMs can be computed from the matching statistics for the `PBWT` as follows. We scan the matching statistics from right to left, and report a SMEM at the column *j* – *A*[*j*].`len` + 1 (of the input matrix) of length *A*[*j*].`len` if either *j* = *w*, or *A*[*j*].`len` ≥ *A*[*j* + 1].`len`. Informally, *A*[*j*].`len` ≥ *A*[*j* + 1].`len` occurs when we cannot extend to the right the current longest common suffix (of length *A*[*j*].`len`) shared by *P* and any row in the input matrix. We show an example of matching statistics for the input matrix `M` in Figure 1.

Next, in Section 3.2, we show how to compute the matching statistics in 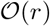-space by storing the following: (1) A mapping structure to support the navigation of the `RLPBWT`; (2) The samples of the prefix array (PA) in correspondence of the beginning and end of each run in the `RLPBWT`; and (3) The *thresholds* identifying the positions of the first minimum divergence array (`DA`) value in each run in the `RLPBWT`. In Section 3.3, we show how to solve Problem 2 in 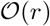-space of a small data structure that we refer to as Φ for the PBWT.

### 3.2 Finding SMEMs in *μ*-PBWT

As previously mentioned, our solution to finding SMEMs in 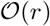-space requires three data structures, which we now describe.

#### 3.2.1 Mapping Structure

Given the position of a bit *σ* in the PBWT, say the *i*-th row and *j*-th column, our mapping data structure returns the positions in the next column of the PBWT of the bits immediately to the right in `M`. This is equivalent to forward stepping in the `PBWT`:

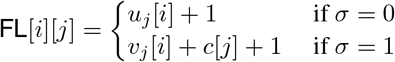

Where i) *u_j_* [i] is the number of zeros until *i* in `col`(`PBWT`)_*j*_ ii) *v_j_*[*i*] is the amount of ones until *i* in `col`(`PBWT`)_*j*_ and iii) *c*[*j*] is the total amount of zeros in `col`(`PBWT`)_*j*_, as in Durbin’s paper.

This mapping allows us to step from one column to the next one (to right) in the `PBWT`. Here, we remind the reader that due to the co-lexicographical ordering on the `PBWT`, it follows that FL-mapping and forward stepping is the analogous counterparts of the LF-mapping and the backward stepping in the BWT. Summarizing, for each column *j* in the `RLPBWT`, we store i) the *r_j_* run head indices *p_j_*, ii) a single r-length data structure *uv_j_* for both *u_j_* and *v_j_*, iii) the integer *c*[*j*], iv) a boolean value *b* storing the symbol of the first run.

In particular, the representation *uv_j_* for both *u_j_* and *v_j_* consists of an interleaved representation for each integer *i*, with 1 ≤ *i* ≤ *r* of the value *v_j_* (or *u_j_*, respectively), up to the start of run *i*, if the *i*-th run consists of zeros (or ones, respectively). For example, given *col*(`PBWT`)_*j*_ = 00101111000000000000 (with *r* = 5), we store: i) *P_j_* = [1, 3, 4, 5, 9], ii) *uv_j_* = [0, 2, 1, 3, 5], iii) *c*[_*j*_] = 15, iv) *b_j_* = ⊤.

#### 3.2.2 PA Samples and Thresholds

Given the `RLPBWT`, we store the positions of the first minimum divergence array (`DA`) value for each run in each column of the `RLPBWT`. We refer to these as *thresholds*. More formally, let `col`(`PBWT`)_*k*_[*i*..*j*] be a maximal run in the *k*-th column of the `PBWT`, we store the `PA` sampled at run boundaries, i.e., the values of `PA`_*k*_[*i*], `PA`_k_[*j*]. We implement the thresholds as bit-compressed integer vectors to store both `PA` samples and thresholds.

#### 3.2.3 Computing the Matching Statistics

Given our data structure, we show how to compute the matching statistics using an algorithm similar to the one used by Rossi et al. [18], which computes the matching statistics in the `BWT`. In particular, we compute the matching statistics in a two-pass algorithm over the input pattern *P*. During the first scan, we process the pattern *P* from left to right, storing for each position the row component of the matching statistics. In the second scan, we process the pattern *P* from right to left, and with the use of a random access data structure on the binary array `M`, we compute the len component of the matching statistics. We assume that we computed the matching statistics component up to position *k* – 1, and are processing the *k*-th column. We let *i* be the row of the `PBWT` that matches the longest suffix of *P*[1..*k* – 1] that is suffix of *S*_1_[1..*k* – 1],…, *S_h_*[1..*k* – 1], and let *p* be the corresponding row in `M` i.e., for all *j* ∈ [1..*h*], `lcs`(*P*[1..*k* – 1], *S*_PA_*k*__,[1..*k* – 1]) ≥ `lcs`(*P*[1..*k* – 1], *S*_PA_*k*_[*j*_[1..*k* – 1]) with *p* = `PA`_*k*_[*i*] where `lcs`(*S, T*) denotes the longest common suffix between two sequences *S* and *T*. Then we distinguish two cases: *match in k-th column*, i.e. when `col`(`PBWT`)_*k*_[*i*] = *P*[*k*] and *mismatch in k-th column*, i.e. when `col`(`PBWT`)_*k*_[*i*] ≠ *P*[*k*]. If we have a match, then row *i* can be used to extend the suffix of *P*[1..*k* – 1] to *P*[1..*k*]; hence we can assign *A*[*k*].`row` = *p*, *A*[*k*].`len` = *A*[*k* – 1].`len` + 1, *i* = `FL`[*i*][*k*], and *p* does not change. Otherwise, if we have a mismatch, it means that for extending the suffix of *P*[1..*k* – 1] to *P*[1..*k*] we need to move to a run before or after the one containing row *i* in `col`(`PBWT`)_*k*_, as the value *P*[*k*] ≠ `col`(`PBWT`)_*k*_[*i*]. Thus let `col`(`PBWT`)_*k*_[*s*..*e*] be a maximal run containing position *i*, then the longest suffix of *P*[1..*k*] that is suffix of *S*_1_[1..*k*], …, *S_h_*[1..*k*] is either the one corresponding to the preceding end or following start of a run of value *P*[*k*] in `col`(`PBWT`_*k*_ with respect to position *i*, i.e., either *S*_PA_*k*[*s*–1_]_[1..*k*] if *s* > 1 or *S*_PA_*k*[*e*+1]__[1..*k*] if *e* < *n*. Since for each run we have stored the samples of `PA` at the beginning and at the end of each run, and we have the value of *p*, we can use the thresholds to decide which candidate to choose. Let *t* be the position of the threshold in the current run. Indeed the thresholds by definition report the positions of the first minimum divergence array (`DA`) value in each run. More precisely, if the position *t* is such that *i* < *t* it means that `lcs`(*P*[1..*k*],*S*_PA_*k*[*s*–1_]_[1..*k*]) ≥ `lcs`(*P*[1..*k*],`S`_PA_*k*[*e*+1]__[1..*k*]) and we can assign *A*[*k*].`row` = *p* = `PA`_*k*_[*s* – 1] and *i* = `FL`[*s* – 1][*k*]. Otherwise, `lcs`(*P*[1..*k*], *S*`PA`_*k*_[_*s*–1_][1..*k*]) ≤ `lcs`(*P*[1..*k*], *S*__PA_*k*_[*e*+1]__[1..*k*]) hence we can assign *A*[*k*].`row` = *p* = PA_k_[*e* + 1] and *i* = `FL`[*e* + 1][*k*].

Once we have collected all the occurrences of maximal matches between the pattern and the matrix, we can compute the lengths of those matches by scanning the pattern *P* from right to left and by comparing the characters in the pattern *P* and in the matrix in correspondence of row *A*[*i*].`row`. To iterate this row, we use the reverse mapping. An illustration of the computation of the matching statistics is shown in Figure 1.

### 3.3 Locating SMEM in *μ*-PBWT

We note that although it is reasonably straightforward to report the number of occurrences of a given SMEM in *S*, it is more challenging to find the location of all the occurrences in *S*. To accomplish this, we store a small data structure that answers queries of the form: given a column index and a prefix array value *j*, return the previous and the next prefix array value in that column. We observe that these two values correspond to rows that we need to consider for finding common suffixes with row *j*—and thus, the occurrence(s) of a SMEM in *S*. We refer to these as Φ-queries in the PBWT.

More formally, given an index *k*, we let `IPA`_*k*_ be the inverse permutation of `PA`_*k*_, i.e. `IPA`_*k*_[`PA`_*k*_[*i*]] = *i*, and define the Φ function [19] for all 1 < *l* ≤ *h* as Φ_*k*_(*l*) = `PA`_*k*_[`IPA`_*k*_[*l*] – 1]. Therefore if `IPA`_*k*_[*l*] = *i*, or equivalently *PA_k_*[*i*] = *l*, it follows that Φ_*k*_(`PA`_*k*_[*i*]) = `PA`_*k*_[*i* – 1], i.e., given a value of `PA`_*k*_ in position *i*, the Φ function returns the preceding value of `PA`_*k*_ in position *i* – 1. Analogously, we can define the inverse of Φ for all 1 ≤ *i* < *h* as 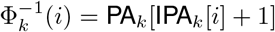. For example, assuming `PA`_6_ = [15, 16, 1, 10, 11, 17, 9, 12, 13, 14, 19, 20, 2, 3, 4, 18, 5, 6, 7, 8] and *i* = 3, we have that Φ_6_(4) = 3 and 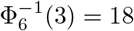. Gagie et al. in [20] showed that the Φ (or Φ^−1^) function for the `BWT` can be stored in 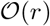 words, and evaluated in 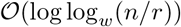-time, where *w* = Ω(log *n*).

To better understand the Φ-function, we observe that whenever we perform an `FL` mapping in a column of the PBWT of two consecutive equal symbols (0 or 1), the resulting positions of the haplotypes in the PBWT are consecutive in the next column after the mapping and their relative order is preserved. Formally, for all 1 ≤ *j* < *w* and for all 1 ≤ *i* < *h*, if `col`(`PBWT`)_*j*_[*i*] = `col`(`pBWT`)_*j*_[*i* – 1] then `FL`[*i*][*j*] = `FL`[*i* – 1][*j*] + 1 = *k* and therefore, `PA`_*j*_[*i*] = `PA`_*j*+1_[*k*] and `PA`_*j*_[*i* – 1] = `PA`_*j*+1_[*k* – 1]. This implies that if we store the `PA` samples at the beginning and at the end of each `PBWT` run, and the whole `PA`_*w*_ column then if we can compute the value of Φ(`PA`_*j*_[*i*])—i.e., compute the value of `PA`_*j*_[*i* – 1]—by performing a `FL` mapping as long as the corresponding `PBWT` values are the same. Now if we assume *k* is the column and *i*′ is the row corresponding to the `PBWT` values mismatch, then we have that `PA`_*k*_[*i*′] = `PA`_*j*_[*i*] and `PA`_*k*_[*i*′ – 1] is sampled since it is at the end of a run. Therefore, by the above observation, we can retrieve the value of `PA`_*j*_[*i* – 1] = `PA`_*k*_[*i*′ – 1]. An example of iterative `FL` mapping to perform Φ queries is depicted in Figure 1d.

We observe that we can use at this point the `DA` samples together with the information of the current row of a SMEM and the next/previous row retrieved by Φ function, to directly check if also the latter shares the same SMEM. Therefore, if we store the `DA` sample at the beginning of each `PBWT` run, while computing the Φ-function for `PA`_*j*_[*i*], we can recover the value of `DA`_*j*_[*i*] as `DA`_*k*_[*i*′] – (*k* – *j*), that is removing from the sampled value `DA`_*k*_[*i*′] the distance travelled by the repeated application of the `FL` mapping.

To avoid performing 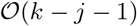 `FL` steps, it is possible to store a successor data structure maintaining the columns where the haplotypes appear as `PA` sample at the beginning of a run and storing the corresponding sample at the end of the previous run and the `DA` sample as satellite information. For example, consider the SMEM in Figure 1 identified by *A*[6].*pos* = 14 and *A*[6].*len* = 6. Since *ϕ*(14) = 13 and `DA`_7_[10] = 6 (having `PA`_7_[10] = 14) then it follows that we know 13-th row shares the same SMEM. Using *ϕ*, we can iterate until *ϕ*(9) = 17, having that `DA`_7_[7] = 1. Using *ϕ*^-1^, we reach 19-th row (*ϕ*^−1^(14) =19) but `DA`_7_[11] = 3, which is less than *A*[6].*len* = 6—so we don not have any other row that share this SMEM.

## 4 Results

We demonstrate the performance of *μ*-`PBWT` by comparing *μ*-`PBWT` with: Durbin’s Algorithm 5 (implemented as `matchIndexed` if the official source code) and Syllable-PBWT [15]. More precisely, for Durbin’s Algorithm 5, we will evaluate a) the memory usage peak and b) the time required for SMEMs finding. For Syllable-PBWT we will evaluate a) the memory usage peak for index construction and b) the size of serialization files. We could not compare *μ*-`PBWT` performance in SMEMS-finding with Syllable-PBWT since Syllable-PBWT implementation allows to computes only L-long matches, that are matches length at least L sites. Therefore, L-long matches are a superset of SMEMs. Finally, we report some statistical results on *μ*-`PBWT`.

### Implementation details

*μ*-`PBWT` is implemented in `C++17` using standard library data structures and relying on the Succinct Data Structure Library (`sdsl`) [21] for succinct data structures implementations such as `int_vectors` and `sd_vectors` with rank and select support. VCF and BCF files input files are supported using the `htslib` library [22].

### Experimental setup

We demonstrate the performance of *μ*-`PBWT` on real-world and simulated datasets. We report the time and memory used for construction and SMEM-locate queries.

We ran experiments on a machine with an Intel Xeon CPU E5-2640 v4 (2.40GHz), 756 GB RAM, and 768 GB of swap, running Ubuntu 20.04.4 LTS (64bit, kernel 5.4.0). The compiler was g++ version 9.4.0 with -`O`3 option. The running time and the maximum resident set size was computed by `/usr/bin/time`.

### Datasets

We first tested *μ*-`PBWT` on all chromosome panels from the 1000 Genome Project. The VCF files were downloaded^1^ and converted to contain only bi-allelic sites via `bcftools view -m2 -M2 -v snps` [23]. The resulting chromosome panels have 5008 haplotypes and a number of bi-allelic sites ranging from ~1 million to ~6 millions. Statistics of the 1000 Genome project panels are in Table 1. Experimentally, we observed these panels are sparse, having indeed fewer ‘1’s compared to ‘0’s. The sparsity of data is confirmed by the average number 11 of runs per column in the run-length encoded PBWT.

**Table 1:** 1000 Genome Project panels information. Columns from left to right report the chromosome number, the number of sites, the average number of runs for each column, the size of the input (in BCF), the size of *μ*-`PBWT` serialization file and the size of Syllable-PBWT serialization file. The last two columns are measured in GB. Each panel has 5008 haplotypes.

| Chr | Sites | Runs | BCF | μ-PBWT | Syllable-PBWT |
| --- | --- | --- | --- | --- | --- |
| 1 | 6 196 151 | 11 | 0.14 | 0.29 | 0.33 |
| 2 | 6 786 300 | 10 | 0.14 | 0.30 | 0.33 |
| 3 | 5 584 397 | 10 | 0.22 | 0.41 | 0.54 |
| 4 | 5 480 936 | 10 | 0.23 | 0.45 | 0.54 |
| 5 | 5 037 955 | 9 | 0.28 | 0.51 | 0.67 |
| 6 | 4 800 101 | 10 | 0.28 | 0.55 | 0.69 |
| 7 | 4 517 734 | 10 | 0.32 | 0.63 | 0.80 |
| 8 | 4 417 368 | 10 | 0.29 | 0.57 | 0.72 |
| 9 | 3 414 848 | 11 | 0.32 | 0.58 | 0.78 |
| 10 | 3 823 786 | 10 | 0.35 | 0.60 | 0.84 |
| 11 | 3 877 543 | 10 | 0.47 | 0.82 | 1.14 |
| 12 | 3 698 099 | 10 | 0.49 | 0.84 | 1.19 |
| 13 | 2 727 881 | 10 | 0.50 | 0.87 | 1.18 |
| 14 | 2 539 149 | 11 | 0.43 | 0.81 | 1.05 |
| 15 | 2 320 474 | 12 | 0.56 | 0.97 | 1.36 |
| 16 | 2 596 072 | 12 | 0.58 | 1.03 | 1.39 |
| 17 | 2 227 080 | 12 | 0.64 | 1.06 | 1.48 |
| 18 | 2 171 378 | 11 | 0.63 | 1.08 | 1.55 |
| 19 | 1 751 878 | 13 | 0.71 | 1.19 | 1.69 |
| 20 | 1 739 315 | 11 | 0.71 | 1.20 | 1.72 |
| 21 | 1 054 447 | 14 | 0.84 | 1.47 | 2.09 |
| 22 | 1 055 454 | 14 | 0.78 | 1.44 | 1.91 |

We used UK Biobank SNP array data across all autosomes (any chromosome that is not a sex chromosome) and high-coverage whole genome sequencing data on chromosome 20 [24]. For the SNP array data, we applied the standard QC recommended by the original authors [24], and phased the data using SHAPEIT4 [25]resulting in 976,754 haplotypes and a total of 670,741 SNPs. For the whole genome sequencing data available on the UK Biobank research analysis platform [1], we used data recently processed and phased by the SHAPEIT5 authors [26], for a total of 300,238 haplotypes and 13,780,193 bi-allelic SNPs and indels on chromosome 20. For the UK Biobank WGS dataset, we applied our method independently to 13 regions of at least 4 megabases and 4 centimorgans on chromosome 20.

Finally, we simulated a 10 megabase region of European samples simulated with msprime [27] with an increasing number of haplotypes up to 2 million (10k, 100k and 1000k individuals, namely panels 1,2 and 3). We also sub sampled the panel with 1000k individuals to obtain two additional panels, namely panels 4 and 5, with the same amount of sites but with 100k and 250k individuals. In Table 2 we have collected the quantitative data of these panels. The BCF files were pre-processed to contain only bi-allelic sites.

**Table 2:** Simulated panels information. Columns from left to right report an ID, the number of sites, the number of haplotypes, the average number of runs for each column, the size of the input (in BCF), the size of *μ*-`PBWT` serialization file and the size of Syllable-PBWT serialization file. The last three columns are measured in GB. In addition, in the last row, we report results on high-coverage whole genome sequencing data on chromosome 20. We were not able to run all the experiments with the Syllable-PBWT due to disk limits, as the input format of Syllabe-PBWT being only uncompressed VCF files.

| Panel | Sites | Haplotypes | Runs | BCF | μ-PBWT | Syllable-PBWT |
| --- | --- | --- | --- | --- | --- | --- |
| 1 | 209 531 | 20k | 9 | 0.04 | 0.06 | 0.25 |
| 2 | 743 171 | 200k | 13 | 0.55 | 0.34 | 8.70 |
| 3 | 2 271 035 | 200k | 4 | 1.2 | 0.49 | - |
| 4 | 2 271 035 | 500k | 8 | 2.6 | 0.78 | - |
| 5 | 2 271 035 | 2000k | 18 | 9.8 | 2.02 | - |
| **Chr20** | **13780193** | **300238** | **-** | **29.6** | **13** | **-** |

### Results on 1000 Genomes Project data

In Figure 2 a) we report: i) memory peak during construction of *μ*-`PBWT` ii) memory peak during querying of *μ*-`PBWT` and iii) Syllable-PBWT memory peak of construction. To improve the readability, we excluded Durbin’s Algorithm 5 memory requirements from the plot, ranging from 60GB to 400GB. We note that our serialization files, as in Table 1 and Figure 4 a), require only twice the memory compared to the input.

**Figure 2:**
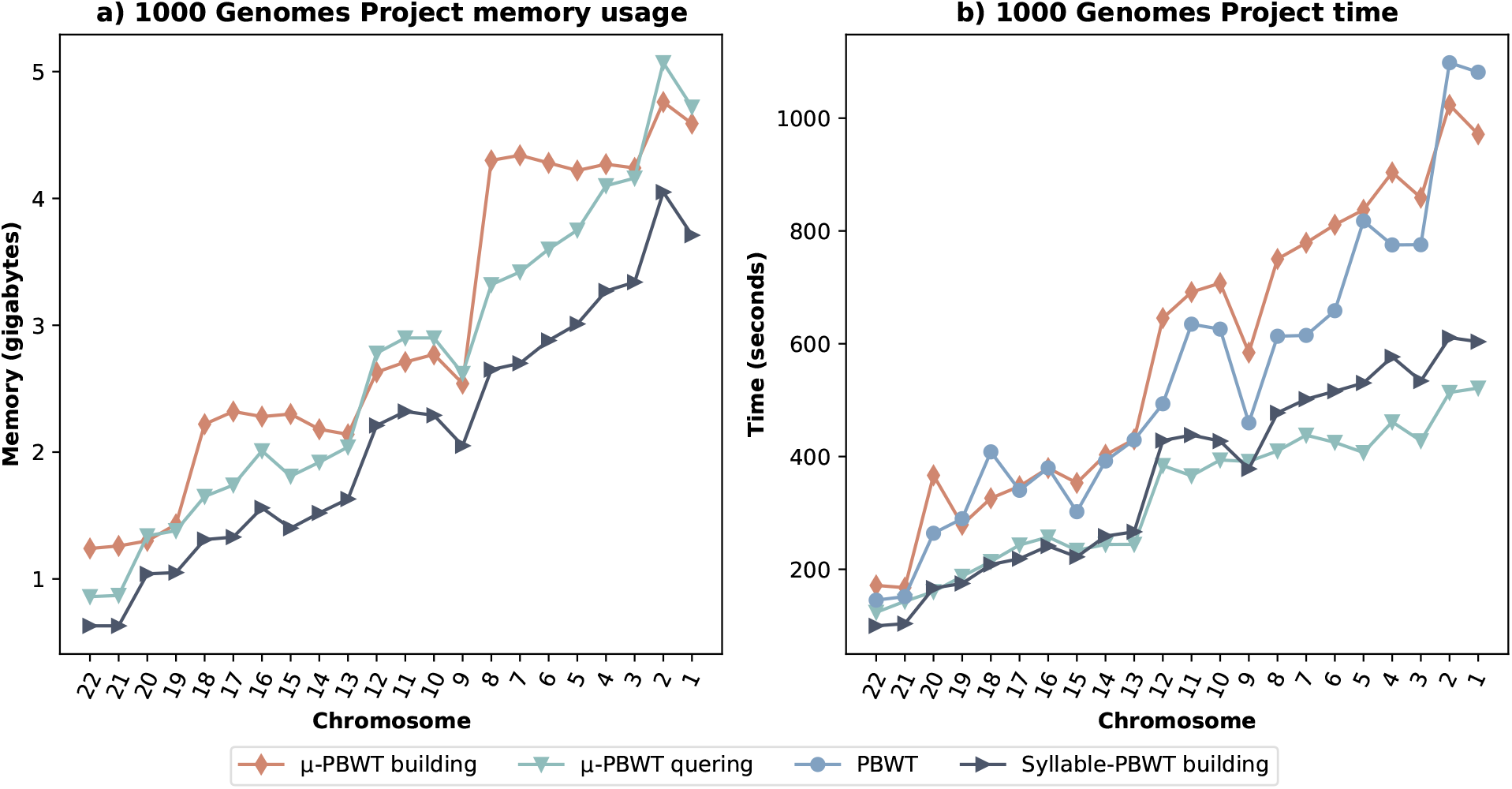
Results comparison on 1000 genome Project data. In a) we have maximum memory usage during building and querying. In b) we have time results, for building, loading and querying.

**Figure 3:**
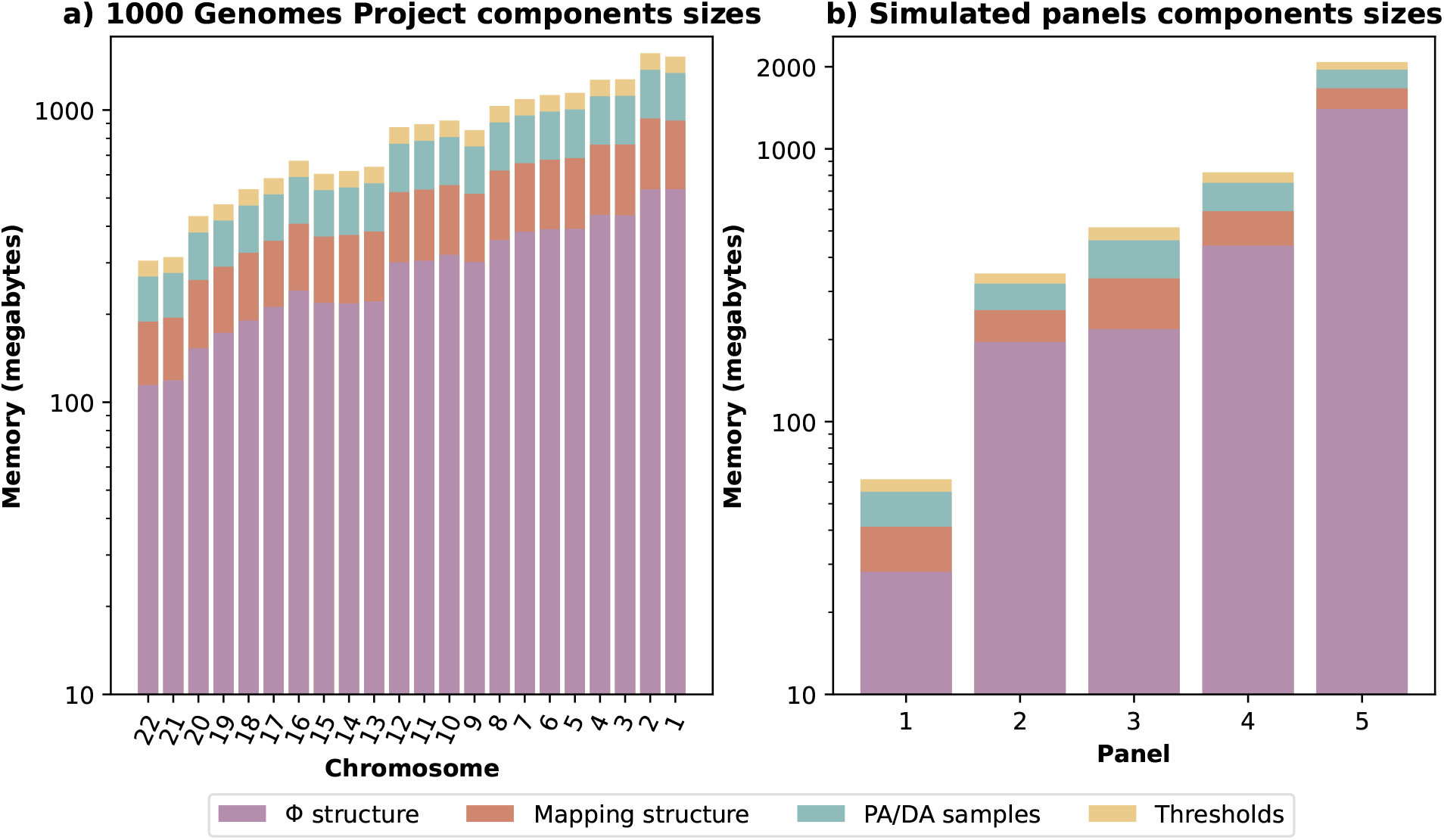
Results comparison on a) 1000 Genome Project data and b) simulated panels, regarding memory usage of the main components of *μ*-`PBWT`.

**Figure 4:**
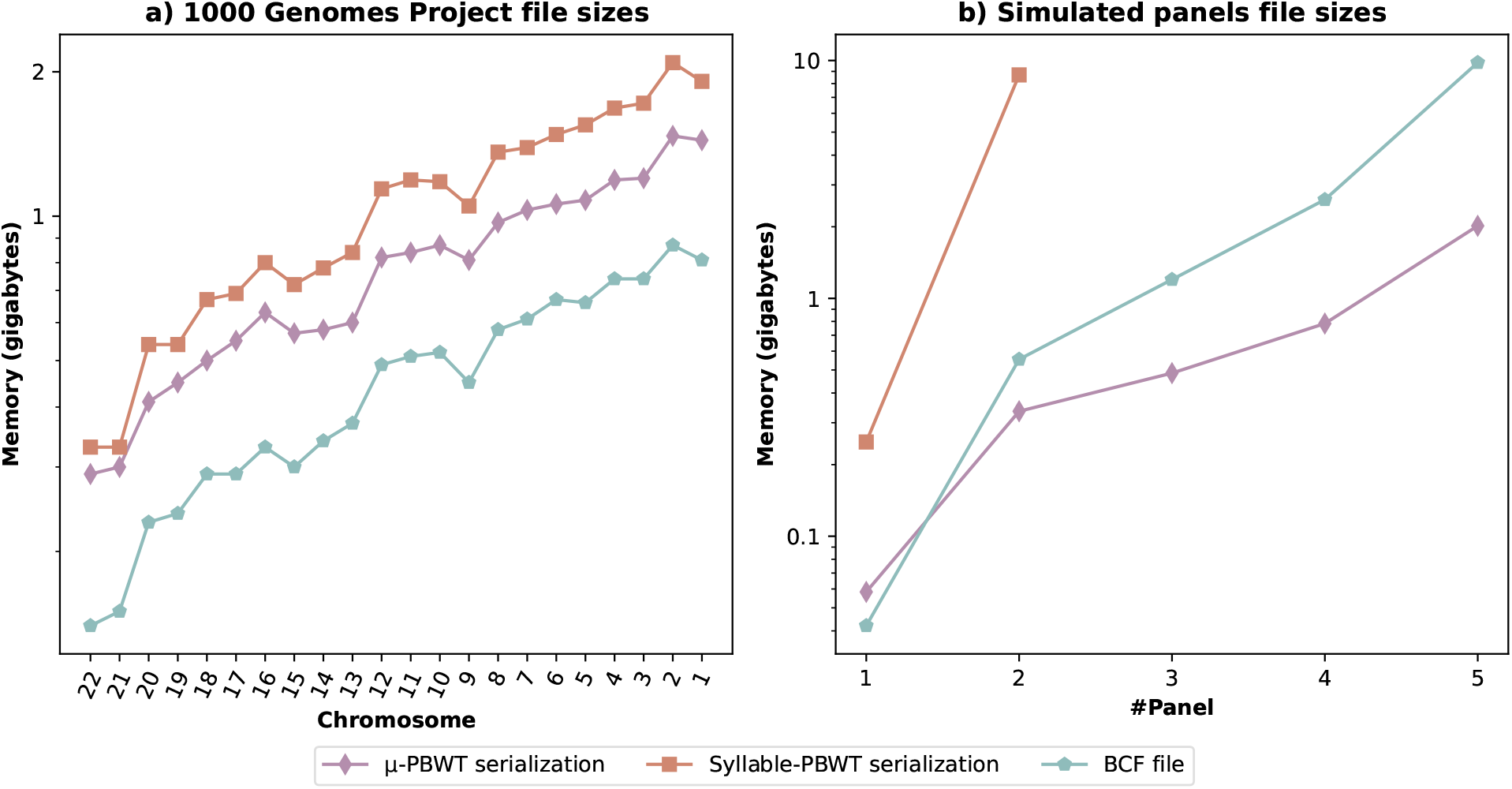
Results comparison on a) 1000 genome Project data and b) simulated panels, regarding BCFs, *μ*-`PBWT` serialization file and Syllable-PBWT serialization files.

Regarding the comparison with Durbin’s Algorithm 5, its memory peak is up to 80 times the memory peak of *μ*-`PBWT` during both building and querying. We have also compared the serialization sizes of Syllable-PBWT. *μ*-`PBWT` requires 25% less memory for the serializations but requires up to twice the memory for building.

To test the performance of computing SMEMs, 100 haplotypes were extracted from the input panels (reduced to 4908 samples), to use them as queries. In Figure 2 b) we can compare the building time with Syllable-PBWT and the SMEM-finding time with Durbin’s Algorithm 5. We note that *μ*-`PBWT` requires twice more time than Syllabe-PBWT for building the index file while regarding the latter our SMEM-finding algorithm requires about twice more time than Durbin’s Algorithm 5 when considering combined building and querying time, since Durbin’s implementation of Algorithm 5 computes most of the necessary arrays at query time. In Figure 3 a) we report the stratification of the memory usage of *μ*-`PBWT` for the mapping structure, PA/DA samples, thresholds, and Φ data structure. The Φ data structure is the component that requires the great amount of memory, since it stores two sparse bitvectors panels and three bit-compressed int-vectors that scale with the total number of runs of the `PBWT`.

### Results on simulated panels and UK Biobank data

We summarize the UK Biobank SNP array data results across all autosomes in Table 3. Due to the low sparsity of these panels, this produced *μ*-`PBWT` with an high number of runs for each column, for example about 13462 for chromosome 20 panel. We run in parallel *μ*-`PBWT` on all the 22 chromosomes, building our serialization files in less than 2 hours. We also applied our method on the UK Biobank high-coverage whole genome sequencing data on chromosome 20. In this setting, our method is able to build an index for the full chromosome 20 in 13 GB of space that represents an almost three times decrease compared to the original gzipped BCF files (stored in a 29GB file), highlighting the potential of our method for compressed genomics on next-generation datasets. Full results are available in Supplementary Material (Table 4).

**Table 3:** Results on UK Biobank SNP array panels. Columns from left to right report the chromosome number, the number of sites, the size of the input (in BCF) and the size of *μ*-`PBWT` serialization file. Each panel has 976754 haplotypes. The last three columns are measured in GB.

| Chr | Sites | BCF | μ-PBWT |
| --- | --- | --- | --- |
| 1 | 54 432 | 4.2 | 13 |
| 2 | 53 433 | 4.2 | 13 |
| 3 | 44 935 | 3.6 | 11 |
| 4 | 41 678 | 3.3 | 10 |
| 5 | 40 020 | 3.2 | 9.6 |
| 6 | 46 515 | 3.9 | 9.4 |
| 7 | 36 682 | 3.0 | 8.9 |
| 8 | 34 141 | 2.7 | 8.2 |
| 9 | 29 581 | 2.4 | 7.7 |
| 10 | 33 086 | 2.7 | 8.2 |
| 11 | 33 827 | 2.7 | 7.8 |
| 12 | 32 011 | 2.6 | 8 |
| 13 | 22 344 | 1.9 | 6.3 |
| 14 | 21 708 | 1.8 | 5.8 |
| 15 | 21 286 | 1.8 | 6.1 |
| 16 | 24 314 | 2.0 | 6.4 |
| 17 | 22 856 | 1.9 | 6.2 |
| 18 | 19 432 | 1.6 | 5.8 |
| 19 | 19 845 | 1.6 | 5.3 |
| 20 | 17 515 | 1.5 | 5.2 |
| 21 | 9940 | 0.8 | 3.4 |
| 22 | 11 160 | 0.9 | 3.6 |

Additionally, more detailed results were obtained on the simulated panels. Only the two smaller datasets (panel 1 and panel 2) allowed experimentation with Syllable-PBWT, which takes as input only raw (not gzipped) VCF files. Syllable-PBWT produced serialization files requiring up to 25 times more space, 16 times more memory and 16 times more time compared to *μ*-`PBWT`. The results are displayed in Figure 4 b). As baseline, we also plot Durbin’s Algorithm 5 estimations on memory usage. On the largest panel, *μ*-`PBWT` reduces the memory consumption of about 25000 times compared to the original PBWT implementation. The average number of runs in each column confirms the high sparsity of these simulated panels, achieving greater effectiveness in the use of run-length encoding and data structures that scale linearly (both in space and time) on the number of runs.

All the serialization files generated by *μ*-`PBWT` are loaded less than 30 seconds on a commodity laptop (AMD Ryzen7 3700U and 16 GB RAM), drastically reducing the hardware requirements for data sharing and analysis whole genome sequencing data.

## 5 Conclusions

In this paper, we present *μ*-`PBWT`, introducing a light index for the PBWT data structure. It leverages the run-length encoding paradigm to solve in small space the problem of finding maximal matches in a set of haplotype sequences. More precisely, we show how significantly it reduces the space requirements for solving two major problems: the SMEMs-finding (i.e. computing maximal matches) and SMEMs-location (i.e. finding occurrences). The main idea behind our method is that *μ*-`PBWT` stores only the information needed to navigate the PBWT by leveraging the runs of haplotypes. Compared to the investigation of the use of the BWT for large genomics data, the PBWT has been comparatively overlooked by the data structures community, even though the increased demand of tools for managing large phased datasets, such as the UK Biobank whole genome sequencing data, for which the PBWT has been originally proposed, making the urgent need of space efficient solutions to store and use these data. Our results address this need, as we show that *μ*-`PBWT` allows a very light indexing of modern biobank data, potentially allowing large whole-genome datasets to be stored and used on a commodity laptop. Results on both the 1000 Genome Project data and simulated panels suggest that *μ*-`PBWT` can scale on whole genome genotype data and it can be used for future analysis on large and repetitive datasets.

**Figure 5:**
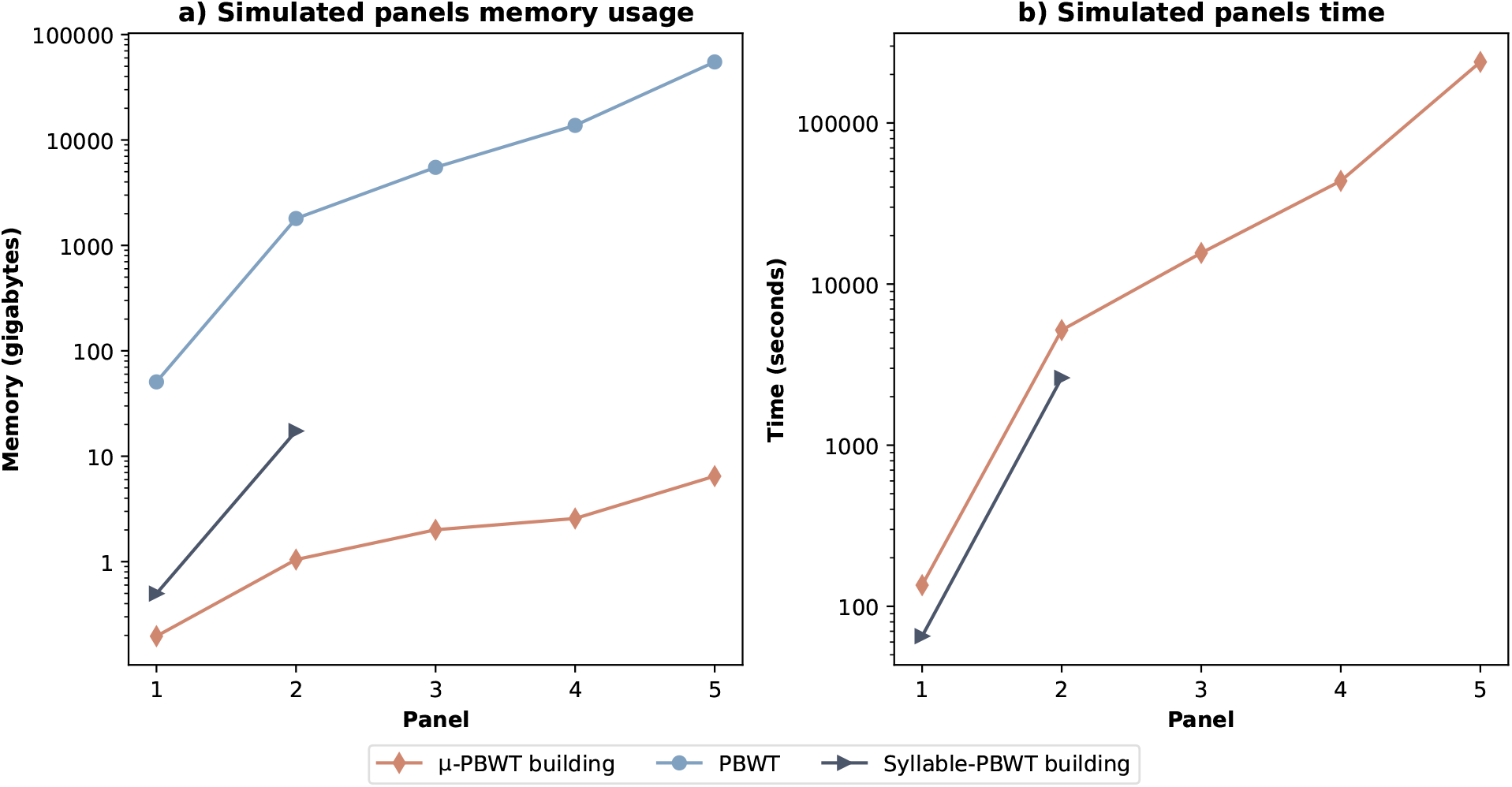
Results comparison on simulated panels. a) Memory usage, combining the size of the various files and the memory peak usages during building. b) Running time for both building and loading the index. Results regarding the original implementation of the PBWT are estimated. We were not able to run all the experiments with the Syllable-PBWT due to disk limits, as the input format of Syllabe-PBWT being only uncompressed VCF files.

## 6 Acknowledgements

The UK Biobank was accessed under UK Biobank project 66995.

## Footnotes

1 Publicly available at https://ftp.1000genomes.ebi.ac.uk/vol1/ftp/release/20130502/

