## Supplemental material for "*μ*-PBWT: Enabling the Storage and Use of UK Biobank Data on a Commodity Laptop"

### Supplementary Material

#### 1000 Genome Project pipeline

The 1000 Genome Project data analysis pipeline has been implemented via Snakemake [1]. The pipeline consists of several steps i) it downloads the 1000 Genome project `vcf.gz` files for chromosomes 1 to 22, ii) it downloads Syllable-PBWT[2], iii) it computes data structure ($\mu$-PBWT[3], Durbin's PBWT[4] and Syllable-PBWT) construction benchmarks, iv) it computes SMEMs finding benchmarks ($\mu$-PBWT and Durbin's PBWT) and v) it makes plots and tables used in this paper. $\mu$-PBWT and Durbin's PBWT are used as Bioconda packages. An overview is available at Figure 6.

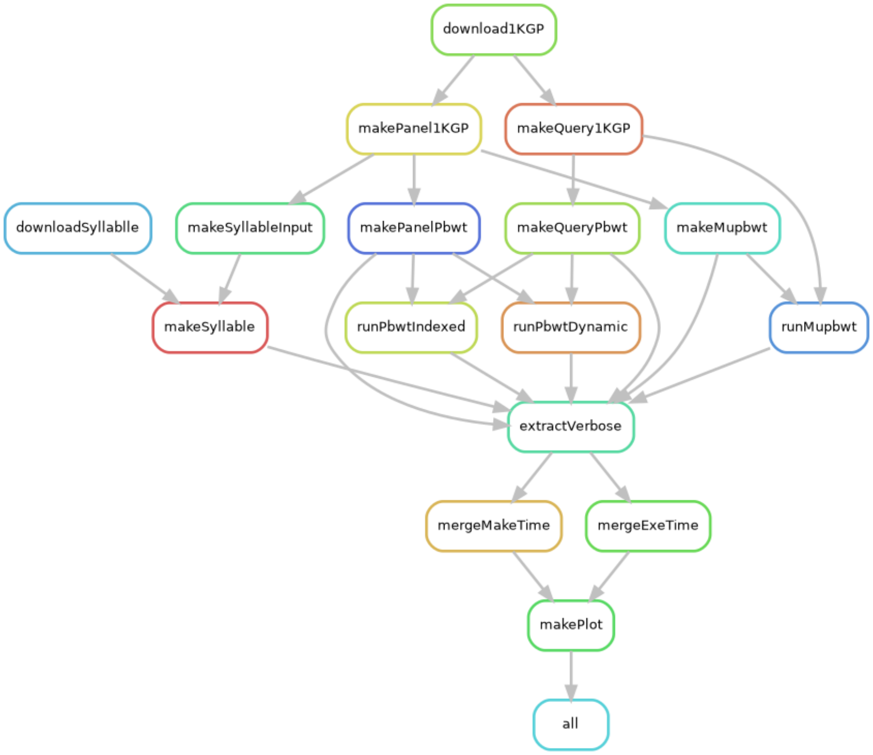


Figure 6: Overview of Snakemake rules for 1000 Genome Project data analysis.

The pipeline is reproducible and it is available at `https://github.com/dlcgold/muPBWT-1KGP-workflow`.

[1]Felix Mölder, Kim Philipp Jablonski, Brice Letcher, Michael B Hall, Christopher H Tomkins-Tinch, Vanessa Sochat, Jan Forster, Soohyun Lee, Sven O Twardziok, Alexander Kanitz, et al. Sustainable data analysis with snakemake. F1000Research, 10, 2021

[2]`https://github.com/ZhiGroup/Syllable-PBWT`

[3]`https://github.com/dlcgold/muPBWT-1KGP-workflow`

[4]`https://github.com/richarddurbin/pbwt`

**UK Biobank**

Table 4: UK Biobank high-coverage whole genome sequencing data on chromosome 20 information. Columns from left to right report the region, the number of samples, the number of sites, the size of the input (in BCF), the size of $\mu$-PBWT serialization file, building time and memory peak usage for construction. In the last row we report the total results. Regarding total build time consider that we have built $\mu$-PBWT for every chromosome 20 region in parallel. The last four columns are measured in GB except for the penultimate which is measured in hh:mm.

| Region | Samples | Sites | Size BCF | $\mu$-PBWT | Time | Memory peak |
|---|---|---|---|---|---|---|
| chr20:60061-4060065 | 150119 | 865267 | 1.9 | 0.88 | 06:25 | 2.27 |
| chr20:4060066-8060066 | 150119 | 880899 | 2 | 0.85 | 06:28 | 2.22 |
| chr20:8060067-12515479 | 150119 | 961591 | 2.1 | 0.77 | 07:04 | 2.05 |
| chr20:12515480-16768988 | 150119 | 917468 | 2 | 0.73 | 06:47 | 1.97 |
| chr20:16768989-21050967 | 150119 | 931010 | 2 | 0.71 | 06:53 | 1.92 |
| chr20:21050968-31549151 | 150119 | 1919134 | 4.2 | 1.20 | 13:54 | 3.06 |
| chr20:31549152-38282825 | 150119 | 1436549 | 2.8 | 0.99 | 10:25 | 2.63 |
| chr20:38282826-43181963 | 150119 | 1056144 | 2.2 | 0.76 | 07:42 | 2.06 |
| chr20:43181964-47619489 | 150119 | 955970 | 2 | 0.79 | 06:56 | 2.09 |
| chr20:47619490-51789198 | 150119 | 923178 | 2 | 0.80 | 06:44 | 2.12 |
| chr20:51789199-55789212 | 150119 | 911452 | 2 | 0.81 | 06:45 | 2.13 |
| chr20:55789213-59874964 | 150119 | 925442 | 2 | 0.84 | 06:49 | 2.20 |
| chr20:59874965-64334101 | 150119 | 1096089 | 2.4 | 0.93 | 08:00 | 2.42 |
| **Total** | **150119** | **13780193** | **29.6** | **11.06** | **-** | **29.15** |
